# Mechanical force regulates the inhibitory function of PD-1

**DOI:** 10.1101/2024.12.13.628463

**Authors:** Hui Chen, Yong Zhang, Lei Cui, Juan Fan, Songfang Wu, Hang Zhou, Yanruo Zhang, Guangtao Song, Ning Jiang, Mingzhao Zhu, Changjie Lou, Wei Chen, Jizhong Lou

## Abstract

The immune checkpoint molecule, programmed cell death 1 (PD-1), plays a pivotal role in regulating T-cell function. Upon binding to its ligands, PD-L1 or PD-L2, PD-1 suppresses T-cell receptor signaling, thereby preventing T-cell activation, making it a critical target in cancer immunotherapy. Although extensively studied, the molecular mechanism of PD-1’s inhibitory function is still not fully understood, especially at the atomic level. Using the biomembrane force probe (BFP), we discovered that interactions between PD-1 and PD-L1/PD-L2 exhibit catch-slip bond behavior under force. At lower forces, catch bonds are observed, transitioning to slip bond as the force increases. Steered molecular dynamics (SMD) simulations revealed a force-induced bond state between PD-1 and its ligands, distinct from the force-free state observed in solved complex structures. Disrupting interactions that stabilize either state weakens the catch bond and diminishes PD-1’s inhibitory function. Interestingly, soluble forms of PD-L1 and PD-L2 compete with their surface-bond counterparts, attenuating PD-1’s suppression of T-cell activation. This suggests that soluble PD-1 ligands could potentially serve as anti-PD-1 drugs. Tumor growth studies in mice confirmed the anti-cancer activity of soluble PD-L1. Our findings highlight the critical role of mechanical force in PD-1’s inhibitory function, suggesting that PD-1 may act as a mechanical sensor in suppressing T-cell activation. These insights indicate that mechanical regulation should be considered when designing PD-1 blocking inhibitors and other PD-1 related cancer immunotherapies.

**Teaser:** This study shows that the interaction between PD-1 and its ligands is force-dependent, and soluble forms of PD-1 ligands can potentially serve as PD-1 blockers in cancer immunotherapy.

## Introduction

Checkpoint immunotherapies targeting co-inhibitory molecules have revolutionized the treatment of various tumors ^1,2^. Among these, programmed cell death 1 (PD-1) stands out as a critical target. Therapies blocking PD-1 or its ligands have achieved remarkable success, illuminating new possibilities for cancer treatment ^3^. Despite several approved drugs for inhibiting PD-1 or its ligands, hundreds more clinical trials are currently ongoing to explore their full potential ^4^. However, a significant challenge remains: many cancer patients exhibit limited response to PD-1 blockade, even for cancers previously shown to be treatable with this therapy ^5^. This limitation underscores the need for a deeper understanding of PD-1’s inhibitory mechanism.

The interactions between PD-1 and its ligands have been explored using various biochemical and biophysical approaches ^6–8^. Antibodies developed against PD-1 or its ligands aim to block PD-1/ligand interactions and diminish PD-1’s inhibitory function ^9,10^. Some of these antibodies have been approved as cancer treatment drugs. However, PD-1/antibody interactions can lead to divergent outcomes ^11^, highlighting the importance of precise mechanisms governing PD-1/ligand interactions and how they correlate with PD-1’s function.

PD-1 is inducibly expressed on T-cell surface post activation. It consists of an extracellular globular domain, a connecting peptide, a transmembrane domain, and an intrinsically disordered intracellular region. Recent evidences indicate that PD-1 can form dimers through transmembrane domain interactions, which linked to its inhibitory function ^3,12^. The intracellular region contains two signaling motifs, ITSM and ITIM, that are essential for PD-1’s activity. Upon binding to its natural ligands, PD-L1 or PD-L2, via the extracellular domain ^13,14^, tyrosine residues within ITSM and ITIM become phosphorylated. This phosphorylation recruits phosphatase SHP-2, which dephosphorylates TCR/CD3, CD28 and other downstream molecules, thereby suppressing T cell activation ^15,16^. Additionally, PD-1 has been shown to inhibit T cell activation by disrupting TCR-CD8 cooperativity ^17^.

Multiple studies confirm that mechanical forces from the intracellular cytoskeleton and intercellular motions influence the formation and context of immunological synapse ^18–20^. These forces inevitably affect the function of the multiple receptor/ligand interactions within the immune synapse. It has been demonstrated that TCR functions of distinguish non-self from self antigens with the aid of mechanical force ^21,22^. Studies have also shown that CD28 co-stimulation could augment CD3 generated traction forces during T cell activation ^23^. Thus, it is reasonable to conclude that PD-1/ligand interactions may also be influenced by the action of force, similar to TCR or CD28. Indeed, studies had confirmed that PD-1 transmits piconewton (pN) forces to its ligands upon surface engagement ^24^. This raises two critical questions: how does mechanical force affect PD-1/ligand interactions affected by the mechanical force, and whether this modulation influences the inhibitory function of PD-1?

In the present study, we used an integrated approach to investigate the interactions between PD1 and its natural ligands. We found that PD-1/PD-L1 and PD-1/PD-L2 interactions exhibit catch bond behavior, where bond lifetimes increase in the low force regime. This force dependence likely promotes PD-1 engagement within the immune synapse, aids in T cell suppression. Given the force dependence of these interactions, soluble and immobilized ligands behave differently on PD-1’s inhibitory function, as soluble ligands can bind PD-1 but unable to provide an external force, and therefore cannot trigger its inhibitory function. We demonstrated that soluble ectodomain of PD-L1 can compete with PD-1 ligands expressed on the surface of APCs for binding PD-1, and serve as a potential PD-1 blocking drug with anti-tumor activity. Conversely, surface-immobilized ligands enable the formation of PD-1 clusters, which protect PD-1 from dephosphorylation and maintain SHP-2 recruitment. This process attenuates T-cell activation by dephosphorylating TCR/CD3 and CD28 in the vicinity of PD-1. Our results suggest that PD-1 may also function as a mechano-sensor. Understanding the mechanical regulation of PD-1 function could provide new avenues for designing PD-1 based immunotherapies for cancer.

## Results

### PD-1 forms catch bond with PD-L1

To investigate whether mechanical force affects the interaction between PD-1 and its ligands, we use biomembrane force probe (BFP) to measure the force dependence of bond lifetimes between PD-1 and PD-L1 at the single molecule level. BFP is particularly suitable for studying interactions between surface receptors and ligands ^21,25,26^. In our experiments, human PD-1 (hPD-1) expressing Jurkat cells or mouse PD-1 (mPD-1) expressing EL-4 cells were repeatedly contacted and detached against beads coated with human or mouse PD-L1 (**Fig. 1A**). We controlled the site density of PD-L1 on the beads to maintain an adhesion frequency below 20%, ensuring predominantly single-bond interactions between PD-L1 and PD-1 on the cell surface (**Fig. S1A and S1B**).

**Figure 1.**
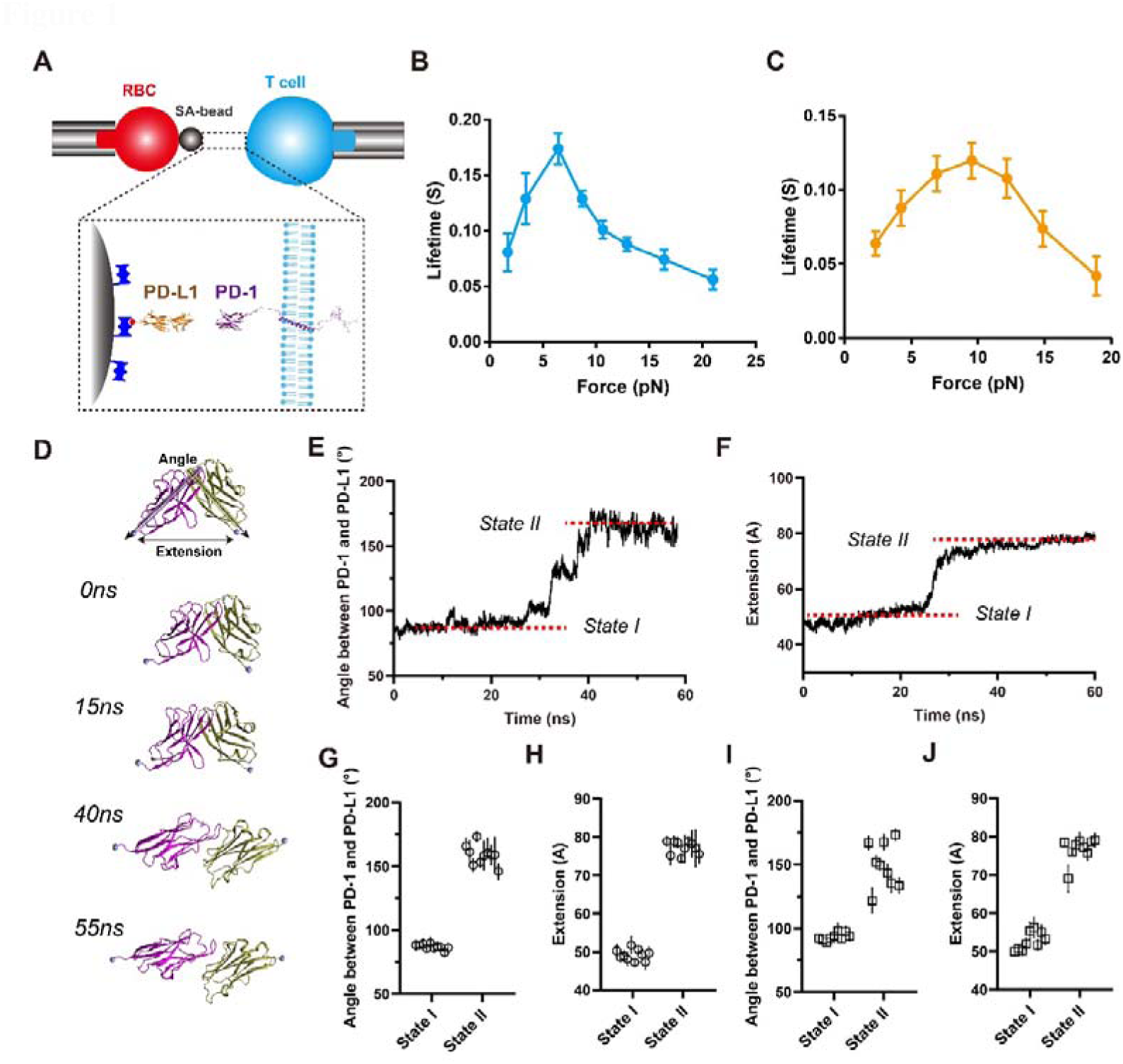
PD-1 forms catch bonds with PD-L1 under force. **A**, Schematic of BFP experiments. PD-1 on a living cell (Jurkat cell for human PD-1 or EL4 cell for mouse PD-1) were driven to interact with human or mouse PD-L1 immobilized on a SA bead; **B-C**, Mean bond lifetime dependence on mechanical force of human (**B**) and mouse (**C**) PD-1/PD-L1 interactions, data are shown as Mean±SEM. **D**, Sequential snapshots at indicated simulation times of cv-SMD simulations showing PD-L1 dissociation from the PD-1; **E**, Representative time-course of the inter-domain angle between PD-1 and PD-L1 in cv-SMD simulations, distinguishing two different binding states; **F**, Representative time-course of the C-terminus to C-terminus (CT-CT) distances between PD-1 and PD-L1 in cv-SMD simulations, showing two different binding states; **G-H**, Statistics of the inter-domain angle (**G**) and CT-CT distance (**H**) between PD-1 and PD-L1 for the two different binding states from ten independent cv-SMD simulations, each point represents an average value of angle on either state in a MD trajectory. **I**-**J**, Statistics of the inter-domain angle (**I**) and CT-CT distance (**J**) between PD-1 and PD-L1 for the two different binding states from nine independent cf-SMD simulations, each point represents an average value of angle on either state in a MD trajectory.

While previous reports suggest that PD-L1 can bind CD80 on T cells ^27^, recent studies increasingly support PD-L1 primarily binding CD80 in cis on the same cell surface ^28–30^. Our data aligns with these findings. Bond lifetimes were collected from PD-1/PD-L1 dissociation events at preset constant force (**Fig. S1C**), revealing that PD-1 forms catch bonds with PD-L1 under low force in both human and mouse systems (**Fig. 1B and Fig. 1C**). Notably, differences can be observed between human and mouse PD-1/PD-L1 dissociation: human PD-1/PD-L1 bonds exhibited longer lifetimes, peaking at approximately 7 pN, compared to mouse PD-1/PD-L1 bonds which peaked at around 10 pN (**Fig. 1B and Fig. 1C**). These results indicate that PD-1/PD-L1 interactions are actually regulated by force, and the force dependent behavior differs between human and mouse.

To elucidate the molecular mechanism of PD-1/PD-L1 catch bonds, we employed steered molecular dynamics (SMD) simulations to model the force-driven dissociation of PD-1 and PD-L1 using both constant-velocity (cv-SMD) and constant-force (cf-SMD) modes. The cv-SMD simulations revealed two distinct binding states (states I and II) during the dissociation process (**Fig. 1D, 1E and 1F**). In state I, the interdomain angles between PD-1 and PD-L1 ectodomains are approximately 87° and the extensions between their C-terminal Cα atoms are around 49 Å, closely resembling the crystal structures (**Fig. 1D, 1G and 1H**). In state II, the interdomain angles are approximately 157° and the extensions are around 77 Å, indicating a novel extended conformation (**Fig. 1D, 1G and 1H**). These two similar binding states were also observed in constant-force SMD simulations (**Fig. 1I-1J and Fig. S1D-S1E**). Interestingly, removing force from state II snapshots reverted the conformation back to state I (**Fig. S1F, S1G**), suggesting reversible conformational changes between states depending on force application. This switching likely enhances the stability of PD-1/PD-L1 interactions under low force, resulting in catch bonds.

### Force-induced PD-1/PD-L1 conformational changes are related to its inhibitory function

To test our hypotheses and further confirm the molecular mechanism of force-regulated PD-1/PD-L1 interactions, we introduced point mutations to PD-1 and/or PD-L1 based on the simulation results.

First, we found that residue E136 of PD-1 can form salt bridges with residues R113 and R125 of PD-L1 in state I, but not in state II (**Fig. 2A-2C and Fig. S2A, S2B**). This interaction thus likely stabilizes state I. This was confirmed by BFP experiments showing that mutations of these residues (R113S or R125S in PD-L1, and E136R in PD-1) weakened the catch bond, while swapping mutations (E136R in PD-1 and R113E or R125E in PD-L1) rescued it (**Fig. 2D-2E**). Co-inhibition assays demonstrated that the mutations weakening the catch bond increased IL-2 secretion, suggesting a reduction in PD-1’s inhibitory function (**Fig. 2F**).

**Figure 2.**
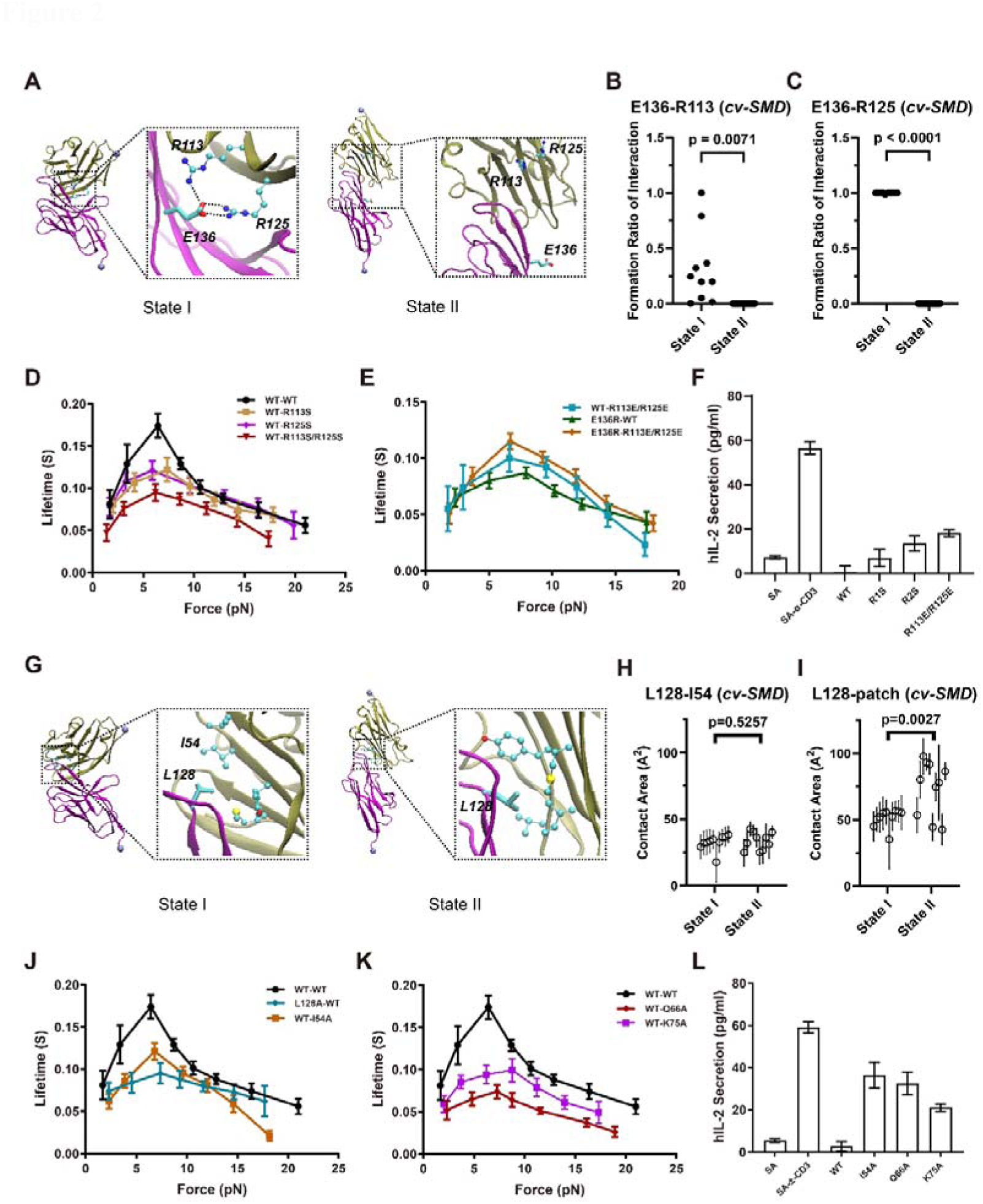
Molecular mechanisms of PD-1/PD-L1 catch bonds. **A**, Representative snapshots showing the interaction between E136 (PD-1) and R113/R125 (PD-L1) in two different binding states; **B** and **C**, Probabilities of salt bridge formation between E136 and R113 (**B**) or R125 (**C**) in cv-SMD simulations (n=10); **D** and **E**, Mean bond lifetime dependence on force of human wildtype or mutated (E136R) PD-1 interacting with human wildtype or mutated PD-L1 (R113S, R125S and R113S/R125S in **D**, R113E/R125E in **E**), data are shown as Mean±SEM. **F**, IL-2 secretion of Jurkat cells expressing human PD-1 stimulated with CD3ε antibody in the presence or absence of widetype or mutated PD-L1; **G**, Representative snapshots showing the interaction between L128 (PD-1) and the hydrophobic patch of PD-L1 in two different binding states as indicated; **H** and **I**, Contact area between L128 (PD-1) and I54 (**H**) or hydrophobic patch (**I**) of PD-L1 in cv-SMD simulations (n=10); **J** and **K**, Mean bond lifetime of wildtype or L128A mutated PD-1 interacting with wildtype or mutated PD-L1, data are shown as Mean±SEM. **L**, PD-L1 mutations impaired PD-1 mediated inhibition of IL-2 secretion.

Next, we observed that residue L128 of PD-1 forms hydrophobic packing with a hydrophobic patch on PD-L1 composed of residues I54, Y56, V68, and M115 in state I, with a contact area approximately 55 Å ^2^. This packing is enhanced in state II, where the contact area is increased to about 80 Å^2^ (**Fig. 2G-2I and Fig. S2C-S2D**). Mutations disrupting this hydrophobic interaction (L128A in PD-1 or I54A in PD-L1) also weakened the catch bond, supporting our hypothesis (**Fig. 2J**). MD Simulations also showed that residue E61 of PD-1 interacts with residue K75 of PD-L1 in state II, but not in state I (**Fig. S2E-S2G**). Additionally, residue Q66 of PD-L1 interacts with different backbone atoms of PD-1 in both states (**Fig. S2H-S2N**). As expected, both Q66A and K75A mutations on PD-L1 also weakened the catch bond (**Fig. 2K**). Co-inhibition experiments confirmed that weakening PD-1/PD-L1 catch bond via these mutations also led to increased IL-2 secretion, indicating a decrease in the inhibitory function of PD-1 on T cell activation (**Fig. 2L**).

To rule out potential interference from protein glycosylation, which has been shown to affect PD1/PD-L1 interactions ^31,32^, we also performed bond lifetime measurements using PD-L1 proteins purified from 293F cells. Although the lifetime values were slightly different, all wild-type and mutant proteins exhibited similar catch/slip bond behavior compared to PD-L1 purified from *E. Coli* (**Fig. S3**). This suggests that glycosylation has minimal impact on PD-1/PD-L1 interactions, comparing to the effects of the introduced the mutations.

Thus, our SMD simulations, single-molecule force spectroscopy (SMFS), and cell functional experiments show that mutations to destabilize either PD-1/PD-L1 binding state impair catch bond behavior and simultaneously reduce PD-1’s inhibitory function.

### PD-L2 forms a weaker catch bond with PD-1, correlating with reduced inhibitory effects

PD-L2, another natural ligand of PD-1, also induces the inhibitory function of PD-1. However, previous studies have demonstrated differences in PD-1 binding to PD-L2 compared to PD-L1^8,33,34^. By measuring the bond lifetime of PD-1 and PD-L2 under force, we found that the interaction between PD-1 and PD-L2 also exhibits catch bond behavior, although it is distinctly weaker than that of PD-1/PD-L1. This weakness is characterized by a lower optimal force and short bond lifetime (**Fig. 3A**). Consistently, the inhibitory function of PD-L2 is also weaker than that of PD-L1 (**Fig. 3B**).

**Figure 3.**
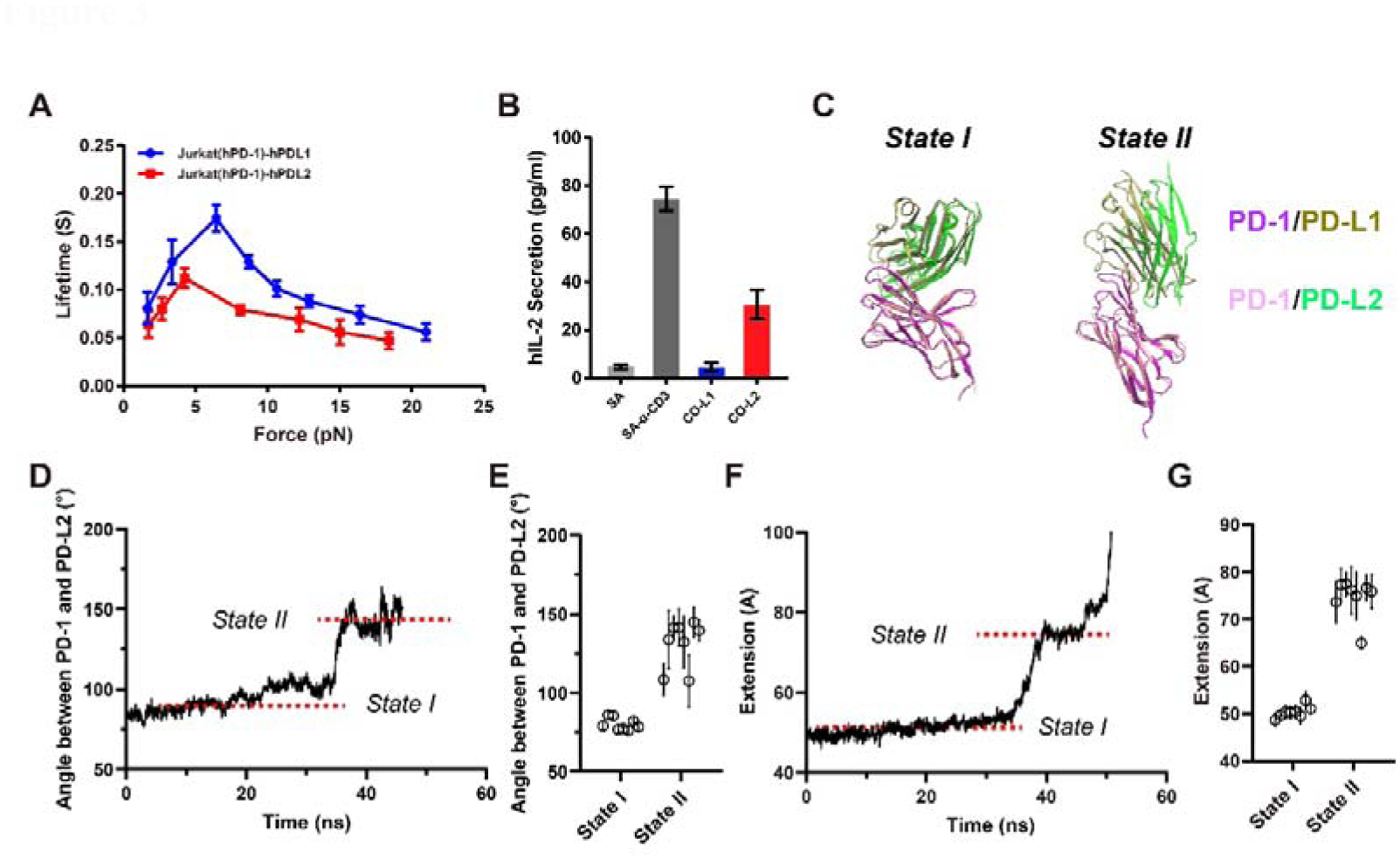
PD-L2 forms weaker catch bonds with PD-1 than PD-L1. **A**, Mean bond lifetime dependence on force for PD-1/PD-L2 interaction, compared to PD-1/PD-L1 interaction, data are shown as Mean±SEM. **B**, IL-2 section inhibition by PD-L1 and PD-L2 in PD-1 expressed Jurkat cells, PD-L1 (CO-L1) or PD-L2 (CO-L2) is coated on SA beads together with CD3ε antibody (SA-α-CD3); **C**, Structural alignment of PD-1/PD-L1 and PD-1/PD-L2 for the two different binding states obtained in SMD simulations; **D**, Representative time-courses of the inter-domain angle between PD-1 and PD-L2 in cv-SMD simulations, distinguishing two different binding states; **E**, Statistics of the inter-domain angle between PD-1 and PD-L2 for the two different binding states in cv-SMD simulations (n=8). **F**, Representative time-courses of the CT-CT distance between PD-1 and PD-L2 in cv-SMD simulations **G**, Statistics the CT-CT distance (**G**) between PD-1 and PD-L2 for two different binding states in cv-SMD simulations (n=8).

As expected, cv-SMD and cf-SMD simulations also revealed two binding states for PD-1/PD-L2 interaction under force (**Fig. 3C-3G and Fig. S4**), similar to the PD-1/PD-L1 complex. Structural alignment showed that PD-1/PD-L1 and PD-1/PD-L2 complexes have similar compact conformation in state I, while exhibit conformational divergence in state II (**Fig. 3C**). In state II, the interdomain angles between PD-1 and PD-L2 are approximately 135°, about 22° smaller than those observed in PD-1/PD-L1 complex (**Fig. 3D-3E and Fig. S4A-S4B**). We propose that the conformational and catch bond differences between state II of PD-1/PD-L1 and PD-1/PD-L2 complexes under force contribute to their functional difference. Specifically, the altered conformation of the PD-1/PD-L2 complex in state II likely result in its weaker catch bond behavior and reduced inhibitory effects.

### Force enhances PD-1 function and its localization at immune synapses

To further understand the role of mechanical force in PD-1 signaling, we constructed a Tension Gauge Tether (TGT) sensor on glass beads ^35^. Biotinylated human CD3 antibodies were tethered to the highest force TGT molecule (56pN), while PD-L1 was ligated to different force TGT molecules (12pN, 23pN and 56pN). We found that the activation of PD-1 signal was highly dependent on mechanical force. The low force TGT molecule (12pN) tethered to PD-L1 could not activate PD-1 signaling and consequently did not attenuate the IL-2 secretion in Jurkat cell (**Fig. 4A and Fig. S5A-S5C**). This result suggests that soluble forms of PD-L1 (S-PDL1) might be ineffective in triggering PD-1 signaling due to their inability to induce mechanical activation of PD-1, a finding supported by previous studies ^36^ and our co-stimulation experiments (**Fig. 4B and Fig. S5D**).

**Figure 4.**
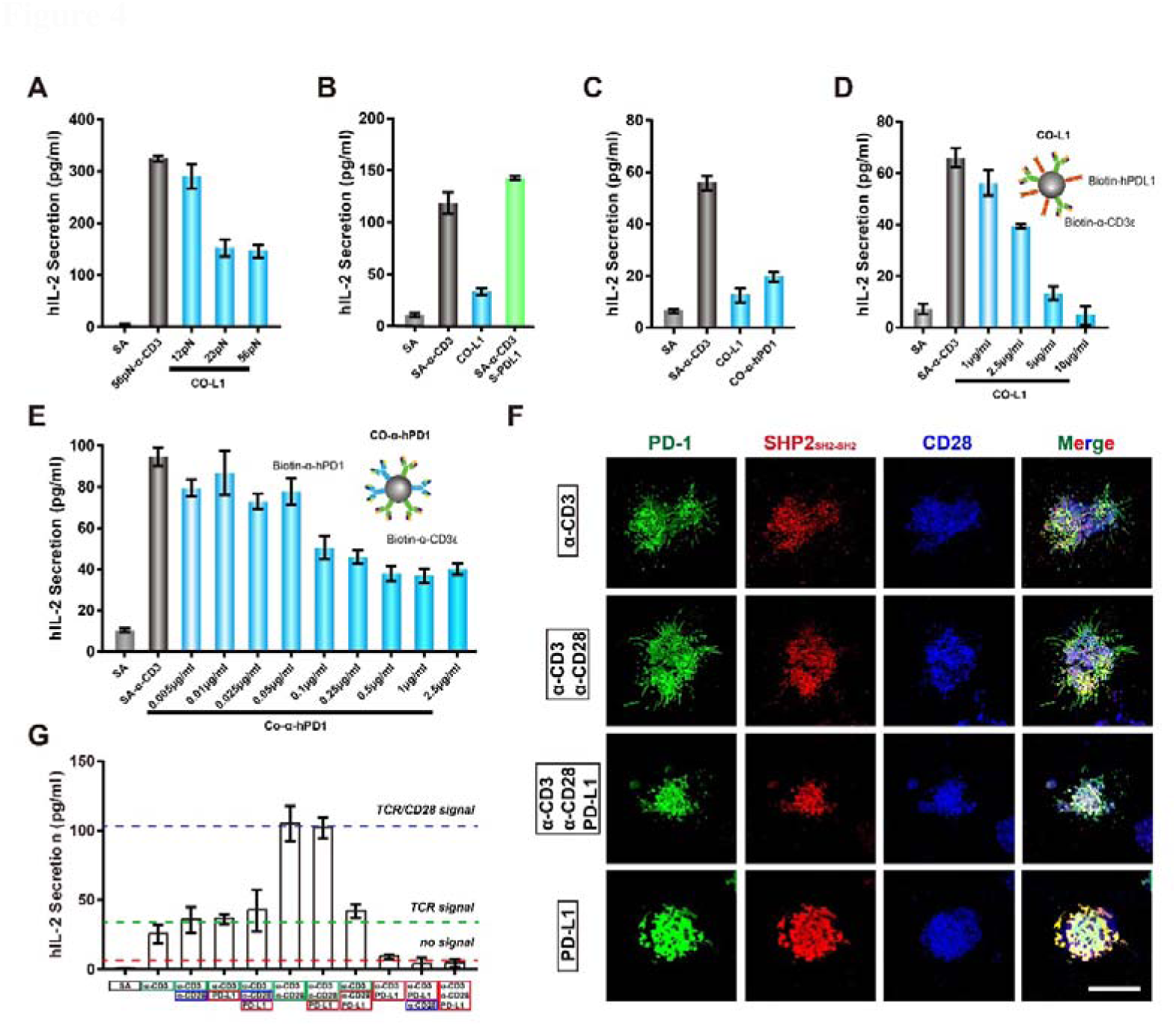
Force enhances PD-1 function and its IS localization. **A**, IL-2 secretion of stimulated Jurkat cells under tension gauge tether (TGT) of indicated tether force; **B-C**, Jurkat cells were stimulated with beads coated with SA-α-CD3, CO-L1, SA-α-CD3 in the presence of S-PDL1 (**B**) or CO-immobilized anti-hPD1 antibody (CO-α-hPD-1) (**C**); **D-E**, IL-2 secretion of Jurkat cells stimulated with anti-hCD3 beads, CO-PDL1(**D**) or CO-α-hPD1 (**E**) at indicated concentration; **F,** PD-1 phosphorylation in Jurkat cells. Jurakt cells (PD-1-mGFP and SHP2_SH2-SH2_-mCherry overexpressed) were stimulated with indicated SLB-ligated proteins for 40 min, Scale bar: 10 μm; **G,** IL-2 secretion of Jurkat cells stimulated with beads coated with indicated proteins. Experiments in A-E were repeated three times in the presence of 2.5 μg/ml anti-CD28 and IL-2 secretion was quantified by ELISA.

To determine whether mechanical force alone is sufficient to trigger PD-1 signaling, we stimulated Jurkat cell with CO-α-hPD1 beads (streptavidin beads simultaneously ligating human PD-1 antibody and CD3 antibody) in the presence of CD28 antibody. The results showed that immobilized PD-1 antibody significantly attenuated IL-2 secretion in Jurkat cells (**Fig. 4C**), aligning with previous studies where immobilized PD-1 antibody inhibited the proliferation of CD4^+^ T cell ^37^.

Unlike PD-L1, which inhibits IL-2 secretion in a dose-dependent manner, PD-1 antibody inhibits Jurkat cell with a constant inhibitory effect at concentrations comparable to those of a CD3 blocking antibody (**Fig. 4D and Fig. S5E**). Considering that the PD-1 antibody has a much higher affinity for PD-1 compared with PD-L1, we investigated the inhibitory function of the PD-1 antibody at lower concentrations. We found that the IL-2 secretion decreased proportionally with increasing of PD-1 antibody concentration (**Fig. 4E**). Additionally, soluble PD-1 antibody could not inhibit IL-2 secretion in Jurkat cell (**Fig. S5F)**.

Thus, we suspect that PD-1 blocking antibody inhibit PD-1 phosphorylation. To monitor PD-1 phosphorylation, we constructed a phosphorylation reporter molecule by tagging the N-terminal SH2 domain of SHP-2 (SHP-2_SH2-SH2_) with mCherry. Stimulating PD-1 overexpressing Jurkat cells with surface-conjugated antibodies revealed that PD-1 could be phosphorylated in the absence of PD-L1, although PD-L1 stimulation alone could result in strongest PD-1 phosphorylation (**Fig. 4F**). These findings are consistent with previous studies indicating that T cell activation without PD-1 stimulation leads to SHP-2 recruitment^15,38^.

Moreover, Successful T cell inhibition requires the co-localization of activated PD-1 with the TCR complex (**Fig. 4G and Fig. S6**) ^16,37^. Studies have shown that elongation of PD-1 ectodomain affects the co-localization with TCR ^38^, potentially explaining why larger antibodies exhibit weaker inhibitory function compared to PD-L1 (**Fig. 4C and Fig. 4E**). Also, a previous study reported that a chimeric receptor, consisting of the mouse CD28 extracellular domain and the human PD-1 cytoplasmic tail, can inhibit CD4^+^ T cell in the presence of co-immobilized mouse CD28 antibody ^15^. Altogether, these findings suggest that mechanical force plays a crucial role in PD-1-mediated T-cell inhibition by regulating the activation and localization of PD-1.

### S-PDL1 serves as a potential agent for PD-1/PD-L1 blockade

The aforementioned results show that S-PDL1 cannot trigger PD-1 signaling. Considering that the density of PD-L1 on the surface might be higher than that in solution, we assessed the function of S-PDL1 in Jurkat cell activation at a higher concentration (200 μg/ml). At this concentration, S-PDL1 slightly inhibited Jurkat cell activation, but the difference was not significant compared to the control group (**Fig. S5D**). This partial inhibition could potentially result from PD-L1 dimerization at high concentration ^39^.

We then co-cultured Jurkat cells with beads coated with both CD3 antibody and PD-L1 (CO-L1) in the presence of S-PDL1 at various concentrations, and found that the IL-2 secretion by Jurkat cell increased with increasing S-PDL1 concentration. Notably, S-PDL1 at 100 μg/ml was sufficient to block the function of immobilized hPD-L1, resulting in IL-2 secretion levels comparable to those stimulated with CD3 antibody alone (**Fig. 5A**).

**Figure 5.**
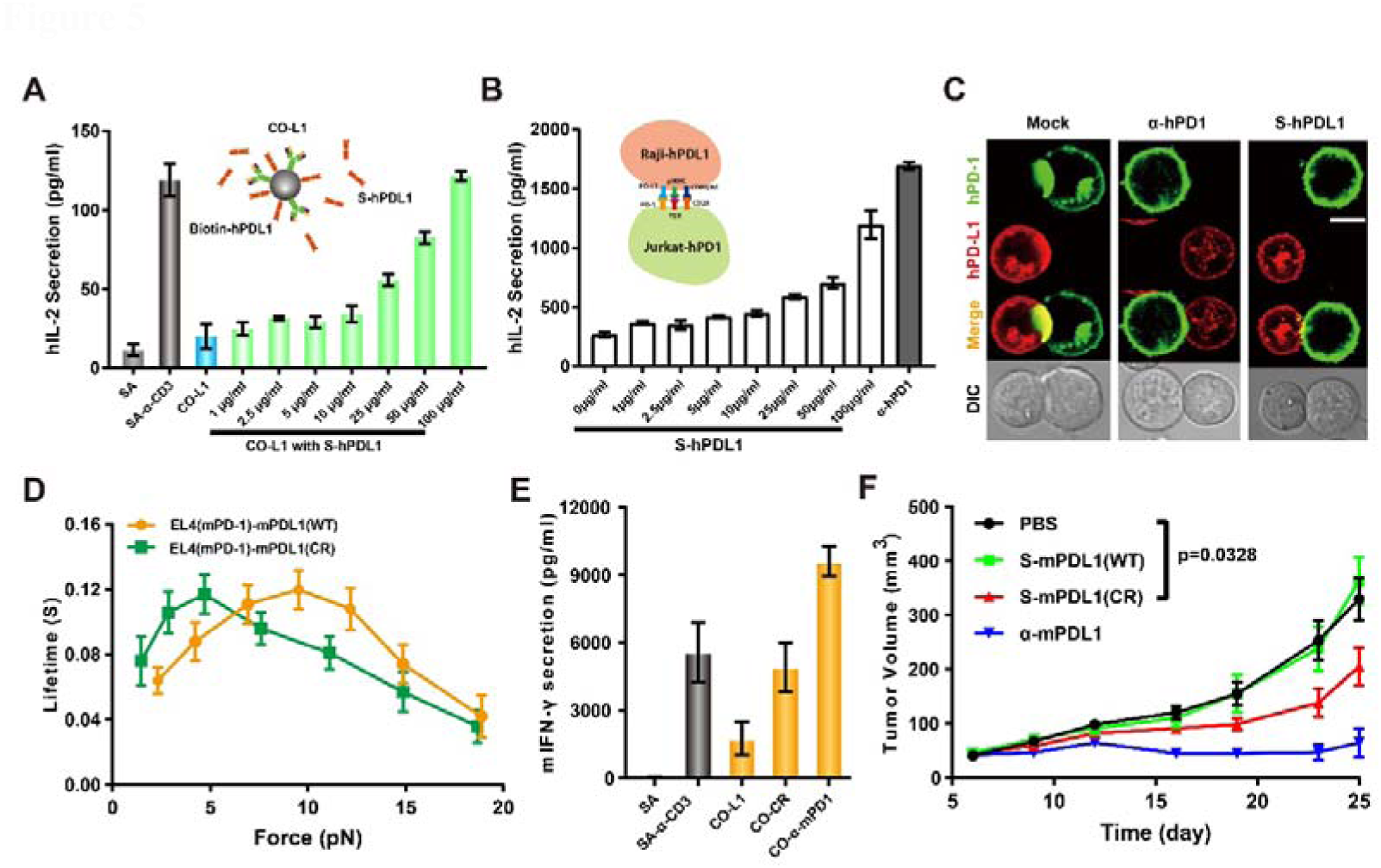
S-PDL1 abrogates PD-1 mediated inhibitory function, similar to PD-1 antibody. **A,** IL-2 secretion of Jurkat cells stimulated with CO-L1 beads in the presence of soluble S-PDL1 at indicated concentration; **B**, IL-2 secretion of Jurkat-hPD1 cells in the presence of S-PDL1 at indicated concentration, 50 μg/ml hPD-1 antibody was used as control. (Inset) A paradigm of hPD-1 expressed Jurkat cells stimulated with hPD-L1 expressed Raji cells; **C**, Representative confocal image of Jurkat (hPD1^+^)-Raji (hPD-L1^+^) conjugates in the presence of S-hPDL1 or α-hPD1, Scale bar: 5 μm; **D**, Force-dependent lifetime of mouse PD-1 interacting with wild type and CR-mutated mPD-L1; **E**, Mouse primary CD8^+^ T cells were stimulated with beads co-immobilized with mPD-L1 or mPD-1 antibody; **F**, Mouse tumor growth Curves, wild type C57BL6 mice were inoculated with MC38 cells and then treated with mouse S-mPDL1(WT), S-mPDL1(CR) or mPDL1 antibody, n=6.

To ensure that the increase in IL-2 secretion was not due to the interactions between PD-L1 and CD80, we detected the surface expression of CD80 on Jurkat cell. The result showed that Jurkat cells do not express CD80, confirming that the increase in IL-2 was not resulted by PDL1-CD80 interaction (**Fig. S5C**). These findings were corroborated by an intact cell assay where Jurkat (hPD1^+^) cells were stimulated with SEE superantigen preloaded Raji (hPD-L1^+^) cells, producing consistent results with the beads-based co-stimulation experiments (**Fig. 5B**). Additionally, a cell-cell conjugate assay showed that S-PDL1 could attenuate hPD-1 aggregation at the immune synapse by blocking the interaction of hPD1 and hPDL1 (**Fig. 5C and Fig. S6B**).

These results indicate the potential antitumor activity of S-PDL1. To confirm this potential and assess the antitumor efficacy of S-PDL1 in vivo, we established MC38 tumors in C57BL6 mice. Considering that wild-type mouse PD-L1 (mPD-L1) has a much lower affinity for mouse PD-1 (mPD-1) in the absence of force loading, we constructed a mouse PD-L1 mutant (mPDL1-CR, a C113R mutation in mouse PD-L1) with a higher binding affinity for mPD-1 (**Fig. 5D and Fig. S7A**). However, mPDL1-CR exhibited no inhibitory functions in either soluble or immobilized form, possibly due to the relatively short lifetime at 10pN (**Fig. 5E and Fig. S7B-S7G**). Surprisingly, the mouse PD-1 antibody also failed to inhibit the activation of CD3^+^CD8^+^ T cells, despite its very high binding affinity for mPD-1 (**Fig. 5E and S7D**). This inconsistency with the Jurkat cell co-stimulation experiments (**Fig. 4C and Fig. 4E**) may be attributed to the differences between human and mouse PD-1 systems.

Importantly, tumor growth curves revealed that soluble mPDL1-CR, but not soluble wild-type mPDL1, could inhibit MC38 tumor growth in C57BL6 mice (**Fig. 5F**). These findings suggest that soluble PD-L1 has potential applications in PD-1 based anti-tumor immunotherapy, and that identifying PD-L1 mutants with higher PD-1 affinity could represent a novel strategy for immunotherapy.

## Discussion

Mechanical force plays a vital role in T cell activation, with the TCR functioning as a mechano-sensor by forming catch bonds with non-self antigens and slip bonds with self antigens ^21,22,40^. In this study, we find that PD-1 can also form catch bonds with PD-L1 and PD-L2 at the single molecule level, identifying key residues on both PD-1 or PD-L1 responsible for these force-regulated interactions. Our MD simulations and SMFS studies reveal subtle differences in the molecular mechanism of PD-1/PD-L1 interactions between human and mouse, suggesting species-specific functions of S-PDL1. TGT-based T cell co-stimulation experiments further demonstrate that mechanical loading on the PD-1/PD-L1 interaction is essential for PD-1 function. A recent paper also described the phenomenon of mechanoregulation of the PD-1/ligand interaction axis and revealed a series of key residues in the PD-1/PD-L2 interaction. These results are in good agreement with our findings ^41^. Collectively, these findings indicate the lifetime of PD-1/ligand bond under force (such as within the immune synapse) is longer than previously thought. This force-enhanced engagement of PD-1 is crucial for its inhibitory functions.

Previous studies showed that PD-1 engagement at the immune synapse is PD-L1 dependent and essential for its inhibitory functions ^38^, with PD-1 capable of transmitting pN forces to its ligand ^24^. This suggests a two-step process, PD-1 is initially recruited to the immune synapse and then stretched by ligand binding for downstream inhibitory function. The stretching induces a conformational change in the PD-1/PD-L1 complex, leading to extended bond lifetime (**Fig. 6**). The prolonged presence of PD-1 allows for its stronger phosphorylation and greater recruitment of SHP-2, which subsequently dephosphorylates CD3 and CD28 in the vicinity of PD-1 (**Fig. 6**). Blocking PD-1 with antibodies or soluble PD-L1 prevents PD-1 recruitment by PD-L1 expressed on APCs ^42^, while mutations in PD-1 or PD-L1 that impair catch bond behavior shorten the dwell time of PD-1 in immune synapse, thereby abrogating its inhibitory function (**Fig. 6**). However, whether PD-1 phosphorylation depends on the force applied by its ligand or antibody binding remains unclear.

**Figure 6.**
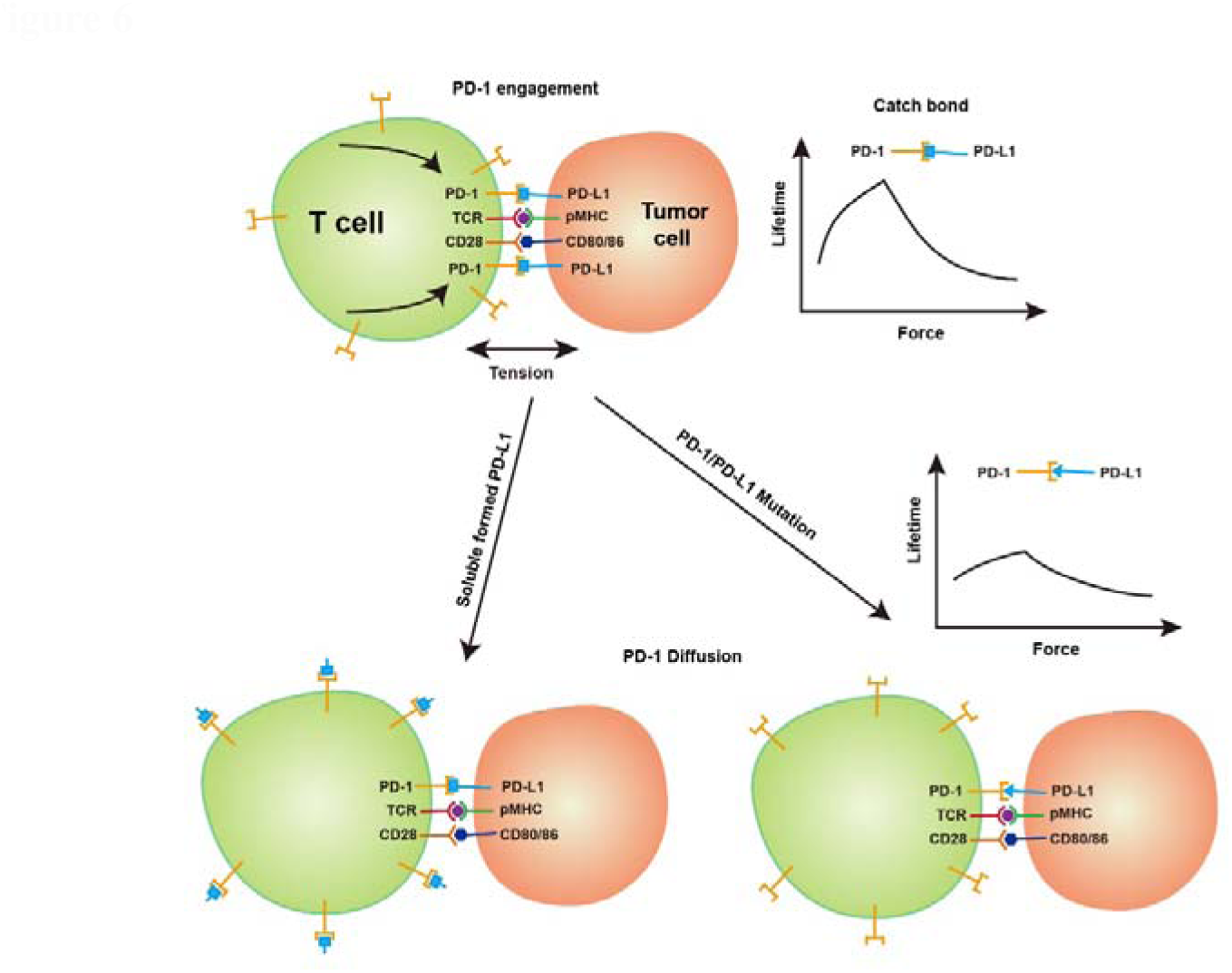
Model of force-regulated PD-1/ligands interaction at the immune synapse. Once the T cell has recognized the target cell via TCR-pMHC interaction, both PD-1 and CD28 bind to its ligand on the target cell side. The resulting tension between the T cell and the target cell acts on the PD1/PDL1 interaction axis, resulting in a conformational change of the PD-1/PD-L1 complex prolongs the retention time of PD-1 in the vicinity of the TCR and CD28, while simultaneously recruiting the downstream SHP-2 to inhibit its signaling. Mutations in PD-1 or PD-L1 result in a weakening of the PD-1/PD-L1 catch bond, which will affect the residency of PD-1 at the immunological synapse, which in turn will impair its inhibitory function. In the microenvironment, free PD-1 ligands can bind to PD-1 on the surface of the T-cell membrane, preventing the aggregation of PD-1 at the center of the immune synapse, blocking PD-1 signaling, and restoring the function of T cells.

Based on previous studies and our current results, we propose two possible mechanisms by which PD-1 induces T cell inhibition. The first, PD-1 on the cell surface may be dynamically phosphorylated or dephosphorylated by Lck or CD45, respectively. Binding with soluble PD-1 ligand does not alter this dynamics balance, but immobilized PD-1 ligand could recruit PD-1 to the cell-cell interface where CD45 is excluded ^43^. The absence of CD45 in the vicinity makes PD-1 more prone to be phosphorylated by Lck. Moreover, PD-1 phosphorylation is also promoted by TCR and CD28 signaling, which facilitates Lck activation ^44^. The second, the intracellular domain (ICD) of PD-1 may adopt in a conformation that protects itself from phosphorylation. Binding with soluble PD-1 ligand can hardly affect this inhibited ICD conformation, whereas surface expressed PD-1 ligand could transmit force to PD-1 and alter this inhibited conformation, which facilitates PD-1 phosphorylation. In both models, the co-localization of PD-1 with TCR and/or CD28 is essential for PD-1 mediated T cell inhibition, and isolated immobilized PD-1 ligands may induce PD-1 phosphorylation but not necessarily T cell inhibition.

Recently, a secreted splicing variant of PD-L1 (sec-PDL1) has been reported to form dimers and inhibit T cell activation ^45^. We speculate that sec-PDL1 could simultaneously bind with CD80 on APCs and PD-1 on T cell, promoting PD-1 engagement on the cell-cell contact interface. Considering that PD-L1 binds CD80 in-cis, soluble sec-PDL1, with its free orientation, can indeed link CD80 and PD-1. Alternatively, sec-PDL1 might induce PD-1 cross-linking via dimerized PD-1, or under the assistance of SHP-2, which has been shown to trigger PD-1 dimerization through its two SH2 domain interacting with two PD-1 molecules ^46^.

The exact mechanism of transmembrane signal transduction of immunoreceptors such as TCR, remains elusive. Mechanotransduction provides a new angle for in-depth investigate ^47^. Our current study highlights the importance of mechanical force in PD-1 signaling by inducing catch bond between PD-1 and its ligands, and soluble PD-L1, where mechanical force application is disabled, could be a potential agent for cancer immunotherapy. However, it remains unclear whether force alone is sufficient to trigger PD-1 phosphorylation. Further studies are required to fully address this question.

## Acknowledgement

We would like to thank Profs. Shengdian Wang and Yan Qin and members in their group for assistance with mouse experiments. The computational resources in this study were provided by the Harbin Supercomputer Center and HPC-Service Station at the Center for Biological Imaging of the Institute of Biophysics. We thank J. Jia and S. Meng (Core Facility, Institute of Biophysics, CAS) for technical support in the flow cytometry analysis.

## Author contributions

J. L. initiated the project, and C. L., W. C. and J. L. conceived the project and designed the experiments. H. C. purified proteins, constructed the cell lines, performed BFP experiments, performed the cell function experiments and analyzed the data. L.C. performed MD simulations and Yong Zhang analyzed the results. J. F participant in protein purification and BFP experiments. H. Z and M. Z. performed mouse tumor studies. S. W., Yanruo Zhang, G. S. helped with the experiments. N. J. provided thoughtful suggestion. H. C. and Y. Z. prepared initial figures and manuscript. J.L., W.C., and C.L. participated in the discussions and optimized figures. All authors helped revise the manuscript.

