## Supporting materials for "Mechanical force regulates the inhibitory function of PD-1"

**Materials and Methods**

**Gene Cloning and Protein Expression, Purification and Labeling**

The extracellular domain of PD-1 and PD-L1 genes were obtained from Generay Biotech and subcloned into pET-28a vector with a C-terminal AVI-Tag. All wild type and mutated proteins for BFP experiments were expressed in Rosetta (DE3) *Escherichia coli* cells (TsingKe, China), purified and refolded according to published protocols ^1^. For wild type and mutated glycosylated PD-L1, genes were subcloned into Phage vector. Recombinant vectors were transduced into 293F cells for protein expression, proteins were then purified using Ni-IDA sefinose resin (BBI life science). The biotinylation of AVI-Tag was performed in vitro using the Biotin-protein ligase kit (GeneCopoeia) and purified by gel filtration chromatography.

**Biomembrane Force Probe (BFP) experiments**

Human red blood cells (RBCs) were biotinylated according to published protocols ^2^. Briefly, fresh human RBCs were isolated from whole blood of healthy volunteers by finger prick. RBCs were covalently linked with biotin-PEG3500-SGA (Jenkem) by incubating at room temperature (RT) for 30 minutes in coating buffer (0.1 M NaHCO_3_, 0.1 M Na_2_CO_3_, pH 8.5, ~180 mOsm). The biotinylated RBCs were then incubated with different concentrations of nystatin in N2 buffer (265.2 mM KCl, 38.8 mM NaCl, 0.94 mM KH_2_PO_4_, 4.74 mM Na_2_HPO_4_, 27 mM sucrose; pH 7.2, 588 mOsm) for 1 hour at 0 ℃, washed twice, and stored in N2 buffer for later BFP experiments.

To prepare protein coated beads, biotinylated proteins were ligated with SA beads (SVP-30-5, Spherotech) by incubating at RT for 30 minutes.

The protein-coated SA bead was attached to the apex of biotinylated RBC aspirated by a stationary micropipette. An hPD-1 expressed Jurkat cell was aspirated by another micropipette driven by a piezoelectric translator. In each typical measurement cycle, the PD-1 expressing cell was brought into contact with the protein-coated bead with a 20 pN impingement force for 0.2 s, and then retracted and held at a desired force to wait for bond dissociation. Bond lifetime was measured from the time of bond sustained at the desired force. The average binned bond lifetime and standard error of the mean (SEM) was plotted as a function of force ^3^.

For adhesion frequency assay, cell was brought into contact with bead for different duration time (t_c_), then retracted till rupture. The adhesion frequency (P_a_) was calculated from the force-time curves.

**Flow cytometry**

For PD-1 binding assay, 1×10^6^ EL4 cells were washed three times with PBS buffer, and incubated with biotinylated WT or mutated mPD-L1 for 30 minutes at 4 ℃. These cells were then further incubated with SA-FITC (eBioscience) for 30 minutes at 4 ℃. After washing with PBS for three times, cells were analyzed on a FACSCalibur (Becton Dickinson) ^4^. Biotinylated BSA and unlabeled mPD-L1 were used as negative control, and biotin-anti-mPD-1 antibody (eBioscience) was used as a positive control.

The density of PD-1 and PD-L1 was quantified using Quantibrite^TM^ PE beads (BD Biosciences) as recommended by the manufacture. Samples were analyzed on a FACSCalibur.

For the quantification of PD-1 expression on Jurkat cells, 1×10^6^ stimulated or unstimulated Jurkat cells were incubated with PE-conjugated hPD-1 antibody (J105, eBioscience) for 30 minutes at 4 ℃. Cells were then washed with PBS and analyzed on a FACSCalibur.

To detect CD80 expression of Jurkat cells, 1×10^6^ Jurkat cells were incubated with biotinylated CD80 antibody (2D10, Biolegend) for 30 minutes at 4 ℃. PBS were used as a negative control. After washing three times with PBS, cells were incubated with SA-APC (eBioscience) for 30 minutes at 4 ℃ and then analyzed on a FACSCalibur. Raji cells were used as a positive control.

**Beads Preparation for Jurkat cell co-stimulation Experiments**

Indicated concentrations of biotinylated anti-hCD3 (UCHT1, Abcam), hPD-L1, hPD-L2, anti-hCD28 (CD28.2, Biolegend) or anti-hPD1 (EH12.2H7, Biolegend) were incubated with SA beads for 30 minutes at RT. These beads were washed three times with PBS and can be directly used for co-stimulated experiments. For preparing CO-beads, the SA-α-CD3 beads were further incubated with serial concentrations of biotinylated PD-L1, PD-L2 or anti-PD1 at RT for 30 minutes. αCD3/αCD28 and αCD3/PD-L1 beads were made by incubating SA-α-CD3 beads with 2.5 μg/ml biotinylated anti-CD28 or PDL1 respectively at RT for 30 minutes. αCD28/PD-L1 beads were made by incubating SA-α-CD28 beads with 15μg/ml biotinylated hPD-L1 at RT for 30 minutes.

**Fabrication of DNA-based tension gauge tether on glass beads**

Glass beads were washed with nanopure water for three times, and then incubated with methanol for 10min. Subsequently, the cleaned beads were suspended with 1% APTES solution (5% H_2_O, 0.5% HAC, 93.5% Methanol) and incubated for 1 h. Silanized glass beads were washed three times with water and then incubated with 2.5% glutaric dialdehyde in water for 1h. The glass beads were incubated with 50 μmol/L NH_2_-labelled ssDNA for 1h. The beads can be stored at 4 ℃ for up to one month. Biotinylated anti-CD3 (or PD-L1) and biotin labelled ssDNA were ligated with streptavidin at a 1:1:1 ratio by incubating for 30 min. Anti-CD3-DNA chimera were annealed with the prepared glass beads by incubating for 40 min. These beads can be used in cell co-culture experiment after three times washing. For PD-1 tension gauge tether assembly, these beads were further incubated with PD-L1-DNA chimera for 40 min. All incubation steps were conducted at room temperature on a rotator to ensure proper mixing and uniform coating.

**Jurkat cell co-stimulation**

For the co-stimulation experiments, 2×10^5^ Jurkat cells were seeded in triplicate wells of a 96-well flat-bottomed plate. Subsequently, 1×10^6^ indicated beads were added to each well, either in the absence or presence of anti-hCD28 (CD28.2, eBioscience). The supernatants from each well were collected at 18-20 hours. The concentration of human IL-2 (hIL-2) secretion was quantified by human IL-2 ELISA kit (Abcam) according to the manufacturer’s instructions ^5,6^.

**Jurakt-hPD1-mGFP stimulation with Raji cells**

For IL-2 secretion assays, Raji-hPDL1-mCherry cells were pre-loaded with 30 ng/ml SEE superantigen (Signalway Antibody) for 1h at 37 ℃. 40 μl of Raji-hPDL1-mCherry cells (5×10^4^ in total) were seeded in a 96-well U-bottom plate and co-cultured with 40 μl Jurkat-hPD1-mGFP cells (2×10^5^ in total). Then 20 μl of either PBS control or various concentration soluble hPD-L1 (s-PDL1) were added into each well. The supernatants were collected at 18-20 hours and quantified by ELISA as described above.

For the Jurkat-Raji conjugates imaging assay, Jurkat-hPD1-mGFP cells and SEE pre-loaded Raji-hPDL1-mCherry cells were plated in an Optical bottom 96-well plate. Soluble hPD-L1 and human PD-1 antibody were added to the solution, corresponding volume of PBS were used as control. Cell conjugates were observed with Olympus FV1000 confocal microscope, and the images were processed using Olympus Fluoview.

**Mouse primary CD8^+^ T cell co-stimulation**

Single cell suspensions were obtained from the lymph nodes of C57BL6 mice according to published protocol ^7^. The cells were stained with anti-mCD3-APC (17A2, eBioscience) and anti-mCD8-FITC (YTS 169AG 101HL, Abcam) at 4 ℃ for 30 minutes. CD3^+^CD8^+^ T cells were then separated by Flow Cytometry (BD FACS Aria IIIu). Beads for mouse primary CD8^+^ T cell co-stimulation were prepared as described above.

For the co-stimulation experiments, 2×10^5^ CD3^+^CD8^+^ T cells and 1×10^6^ indicated beads were incubated at 37 ℃, 5% CO_2_ for three days in the absence or presence of anti-mCD28(37.51, eBioscience). Supernatants were then collected for the quantification of IFN-γ secretion using an ELISA kit (eBioscience). To detect PD-1 expression, the stimulated cells were firstly incubated with APC-mPD-1 followed by PBS washing, and then quantified by Flow-cytometry (Becton Dickinson).

For cell proliferation assays, cells were labeled with 1 μM CFSE (eBioscience) for 10 minutes at RT in the dark. The labeled cells were further co-cultured with indicated beads for three days, and subsequently analyzed by Flow-cytometry.

**TIRF-SIM**

Supporting lipid bilayers (SLBs) consisting of 95% DOPC, 5% DSPE-PEG-biotin, 0.1% DSPE-PEG5000 were prepared as previously described ^8^. Biotinylated CD3ε and CD28 antibodies and PD-L1 were attached to the SLB via streptavidin. Jurkat cells expressing PD1-mGFP and CD28-mCherry were added to the SLB and incubated at 37 ℃ for 40 min. After incubation, cells were fixed with paraformaldehyde (PFA), and stained with APC-labeled Lck antibody. Imaging was performed using a Multi-SIM microscopy equipped with 100× oil immersion objective ^9^. Image processing and data analyses were carried out using ImageJ.

**Tumor growth and treatment**

2×10^5^ MC38 tumor cells were injected subcutaneously into the flank of C57BL/6 mice. 50 μl (1 mg/ml) PD-L1, PD-L1-CR, αPD-L1 mAb or PBS were injected intratumorally at days 6, 9, 12 and 16 post-tumor inoculation. Tumor volumes were measured using a caliper and calculated according to the formula: length ×width ×height/2 as previously described ^10^.

**Molecular dynamics (MD) simulation**

The complex structures of the human PD1 Ig-like V-type (IgV) domain complexed with PD-L1 IgV domain, and complexed with PD-L2 IgV domain, were built using the PDB structures (PDB code: 4ZQK^11^ & 6UMT ^12^) as initial models, respectively. The missing residues of PD-1 (Asp85-Asp92) and PD-L2 (D65-S67) were built using the Modloop server ^13^. The protein complexes were solvated in a rectangular box of TIP3P water molecules. Na^+^ and Cl^-^ ions were added to neutralize the whole systems (~0.15 M). Both systems were processed using the VMD program and the CHARM36m force field for proteins ^14,15^.

The resulting systems were first pre-equilibrated to relax the added missing region of protein, counter ions and water molecules. Subsequently, the production simulations lasted ~100 ns with 2fs time-step. Four independent repeating simulations were performed. During these simulations, the temperature of the systems was maintained at 310 K with Langevin dynamics, and the pressure was controlled at 1 atm with the Nosé-Hoover Langevin piston method. Particle Mesh Ewald summation was used for electrostatic calculation and a 12 Å cutoff was used for short-range non-bounded interactions. The simulation trajectories were recorded every 20 ps.

Representative snapshots of each production run for the PD-1/PD-L1 and PD-1/PD-L2 complex systems were extracted as initial conformations for steered molecular dynamics (SMD) simulations. Before the forces were applied, these snapshots were first simulated under temperature and pressure control with 1fs time-step for 1 ns and underwent free dynamics simulations for another 1 ns for relaxation. The final confirguations were used for the following SMD simulaitons.

Both constant-velocity (cv-SMD) and constant-force SMD (cf-SMD) simulations were performed to investigate the dissociation process of PD-1 and PD-L1/L2. In each cv-SMD simulation, the C-terminal Cα atom of the PD-1 IgV domain was constrained at its initial position with a spring of spring constant ~1400pN/nm, and the C-terminal Cα atom of PD-L1/L2 IgV domain was pulled with a dummy spring of spring constant ~70pN/nm which moved at a speed ~0.1 nm/ns s. In cf-SMD simulations, the applied force was held at 100pN or 50pN until the complex dissociated. For each system, multiple SMD trajecties were obtained for statistical analysis. The Energy minimizations and MD simulations in this study were performed with NAMD ^16^ under periodic boundary conditions.

The interdomain angle between PD-1 and PD-L1/L2 was used to depict their relative orientation, which was defined as the angle between the vector on PD-1 residues (a vector connecting centroids of S57-S62, L100-R104, L128-A132 and T45-N49, S71-Q75, V111-S118 backbone atoms) and that on PD-L1/L2 (a vector connecting centroids of F42-L53, Q91-G95, Y118-D122 and E31-N35, S80-R84, I101-A109 backbone atoms for PD-L1; of D45-I55, L80-G83, Y106-A109 and E33-N37, E63-S67, P90-G98 backbone atoms for PD-L2). The formation of salt bridge interaction was defined as a distance smaller than 3.5 Å between the heavy atoms of the donor and the acceptor residue.

**Statistical Analysis**

All statistical analyses were performed using GraphPad Prism. Unless otherwise stated, data were presented as means ± SD or scatter plots. Calculated P values were reported. Differences between the means of experimental groups were calculated using Student’s t test. Comparison between tumor sizes was done using two-way ANOVA.

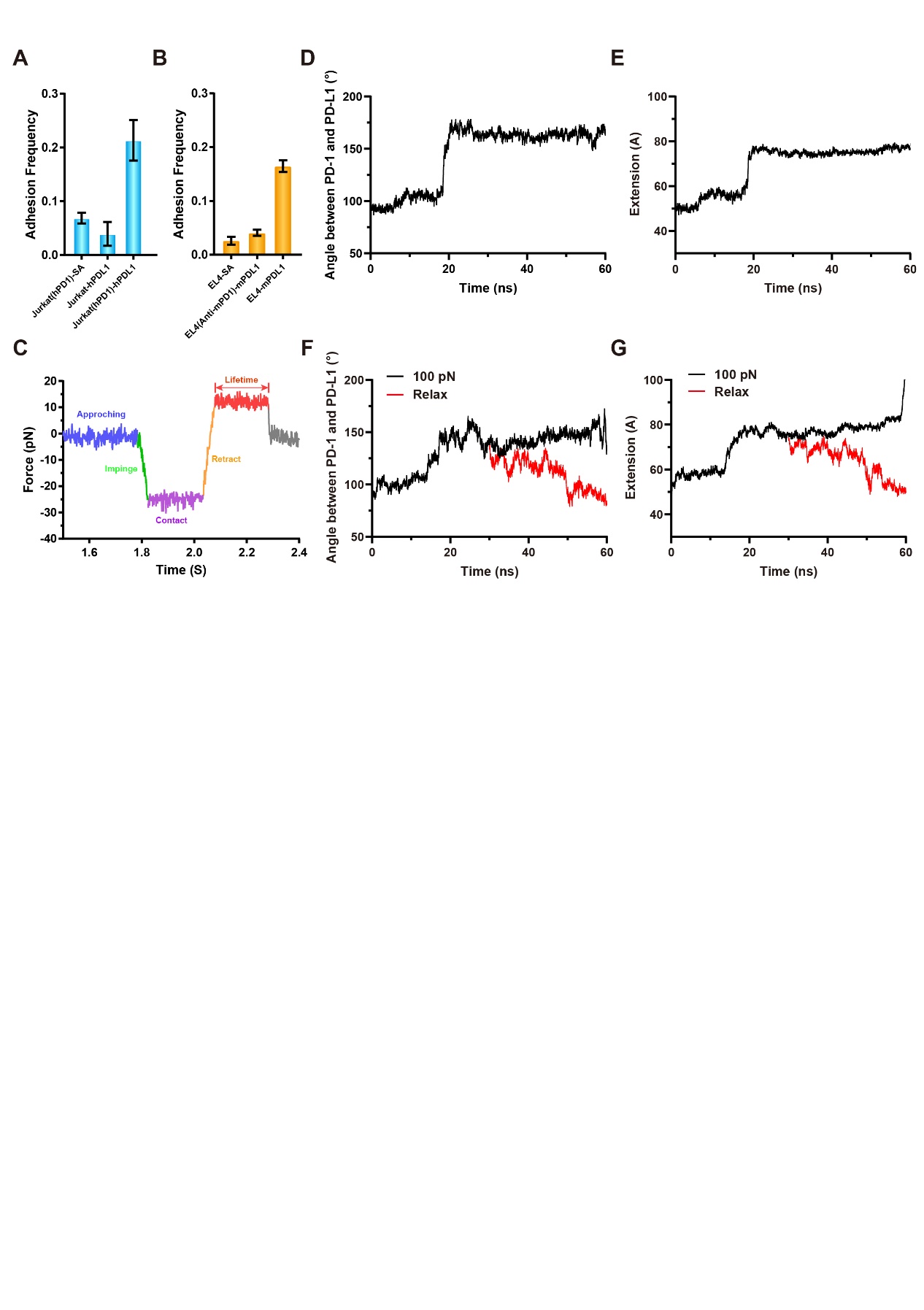

**Figure S1. Characterization of PD1/ligands interactions.**

**A-B**. The adhesion frequency of human (**A**) and mouse (**B**) PD-1/PD-L1 interactions, compared to the indicated control, data are shown as Mean±SD.

**C**. Typical force-clamp curve from BFP experiments, different phases were shown in differed colors and indicated;

**D**-**E**. Representative time-course of the inter-domain angle (**D**) and CT-CT distance (**E**) between PD-1 and PD-L2 in cf-SMD simulations.

**F**-**G**. Representative time-course of the inter-domain angle (**F**) and CT-CT distance (**G**) between PD-1 and PD-L2 in cf-SMD simulations (black) and relax simulations (red).

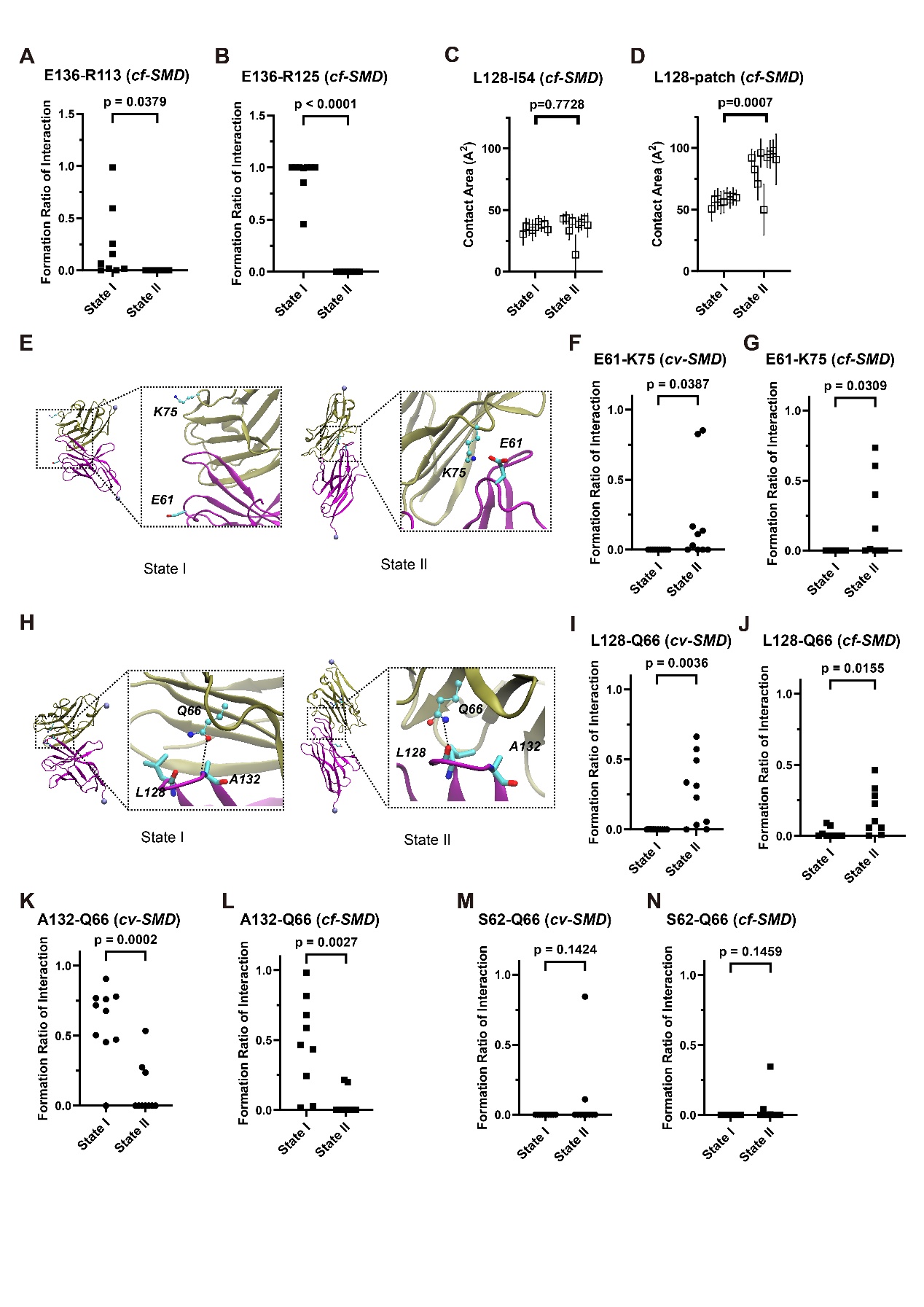

**Figure S2. Key residue pairs identified from MD simulaitons.**

**A-B**. Probabilities of the formation of E136/R113 (**A**) and E136/R125 (**B**) from PD-1 and PD-L1 in cf-SMD simulaitons (n=9).

**C-D**. Analysis of contact area between L128 (PD-1) and I54 (**C**) or hydrophobic patch (**D**) of PD-L1 in cf-SMD simulations (n=9);

**E**. Representative snapshots showing the interaction between E61 (PD-1) and K75 (PD-L1) in the two different binding states.

**F-G**. Probabilities of salt bridge formation between E61 (PD-1) and K75 (PD-L1) in cv-SMD simulations (**F**, n=10) and cf-SMD simulations (**G**, n=9).

**H**. Representative snapshots depicting interaction between L128/A132 backbone (PD1) and Q66 sidechain (PD-L1) in the two different binding states.

**I-J**. Probabilities of hydrogen bond formation between L128 backbone and Q66 sidechain in cv-SMD simulations (**I**, n=10) and cf-SMD simulations (**J**, n=9).

**K-L**. Probabilities of hydrogen bond formation between A132 backbone and Q66 sidechain in cv-SMD simulations (**K**, n=10) and cf-SMD simulations (**L**, n=9)..

**M-N**. Probabilities of salt bridge formation of S62/Q66 in cv-SMD simulations (**M**, n=10) and cf-SMD simulations (**N**, n=9).

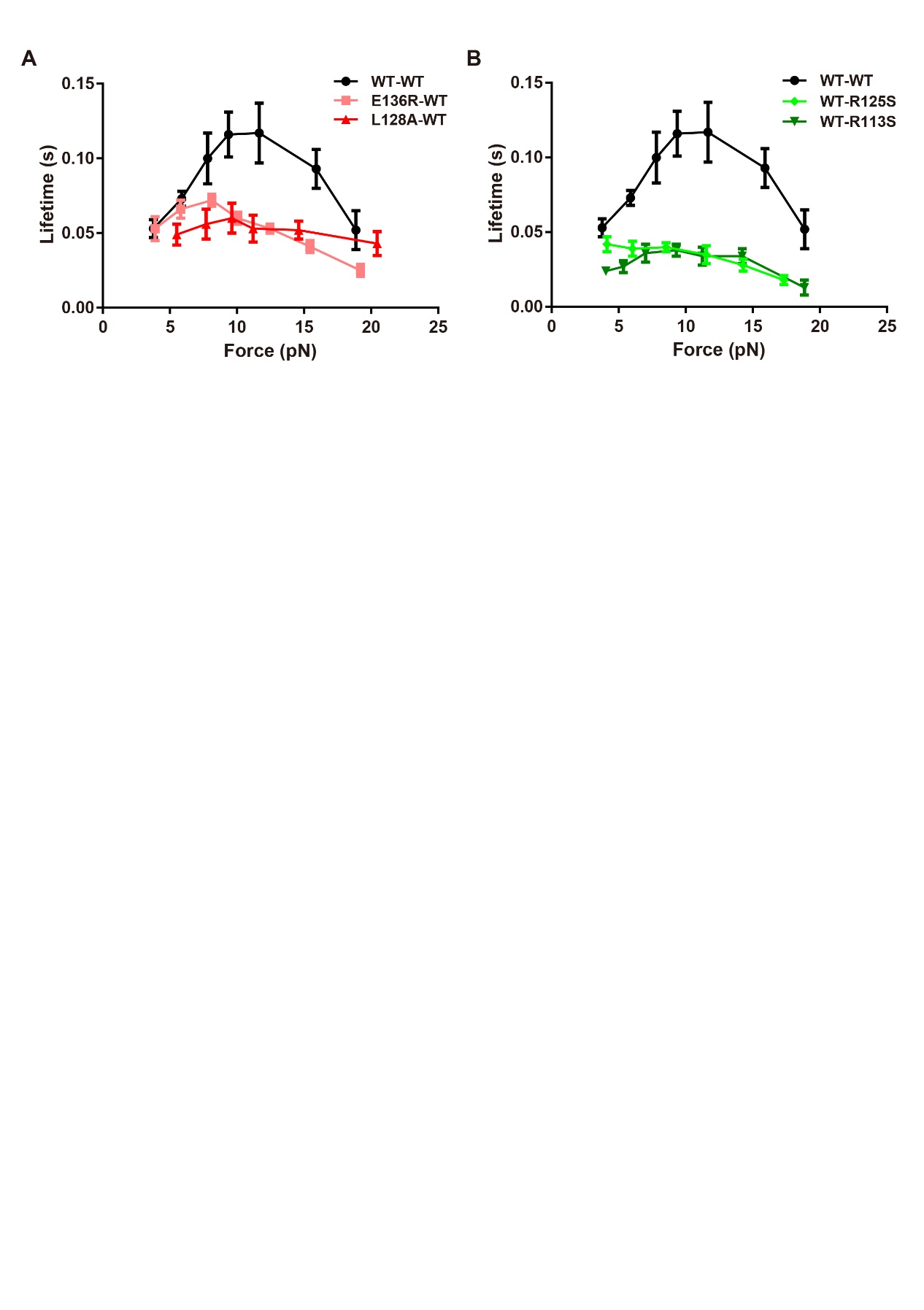

**Figure S3. Protein glycosylation does not alter catch bond behavior of PD-1/PD-L1 under force.**

(**A**) Mean bond lifetime dependence on force for wt or mutated (as indicated) human PD-1 interacting with wt human PD-L1.

(**B**) Mean bond lifetime dependence on force for wt PD-1 interacting with wt or mutated (as indicated) PD-L1.

Indicated wt or mutated proteins were expressed and purified from 293F, data are shown as Mean±SEM.

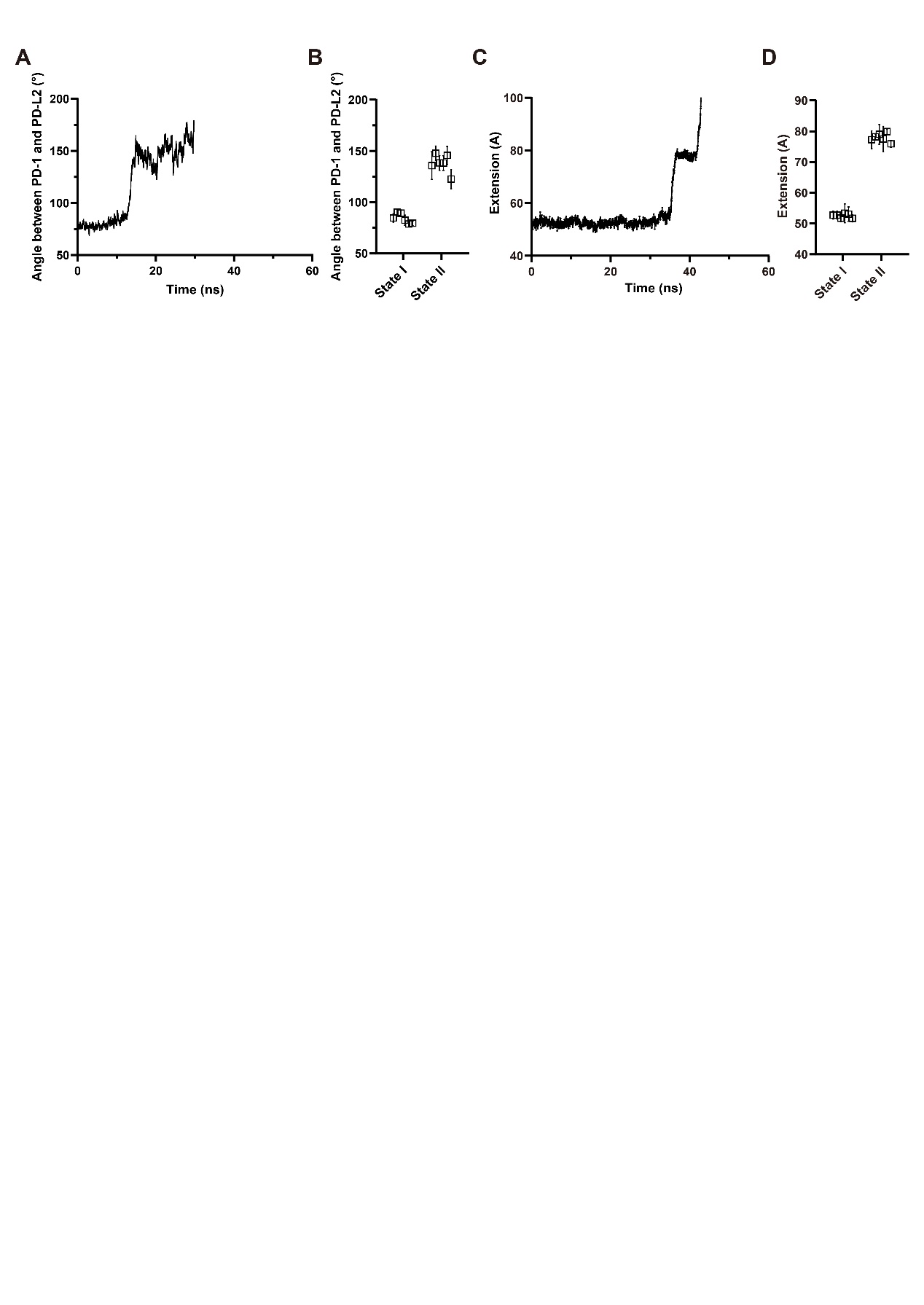

**Figure S4. PD-1/PD-L2 dissociation by cf-SMD simulations.**

**A.** Time-course of the inter-domain angle between PD1 and PD-L2 in one representative cf-SMD simulations, exhibiting two different bending sates.

**B.** Statistics of inter-domain angle between PD-1 and PD-L2 for the two binding states in cf-SMD simulations (n=6).

**C.** Time-course of the CT-CT distance between PD-1 and PD-L2 in the representative cf-SMD simulations shown in A.

**D.** Statistics of CT-CT distance between PD-1 and PD-L2 for the two different binding states in cf-SMD simulations (n=6).

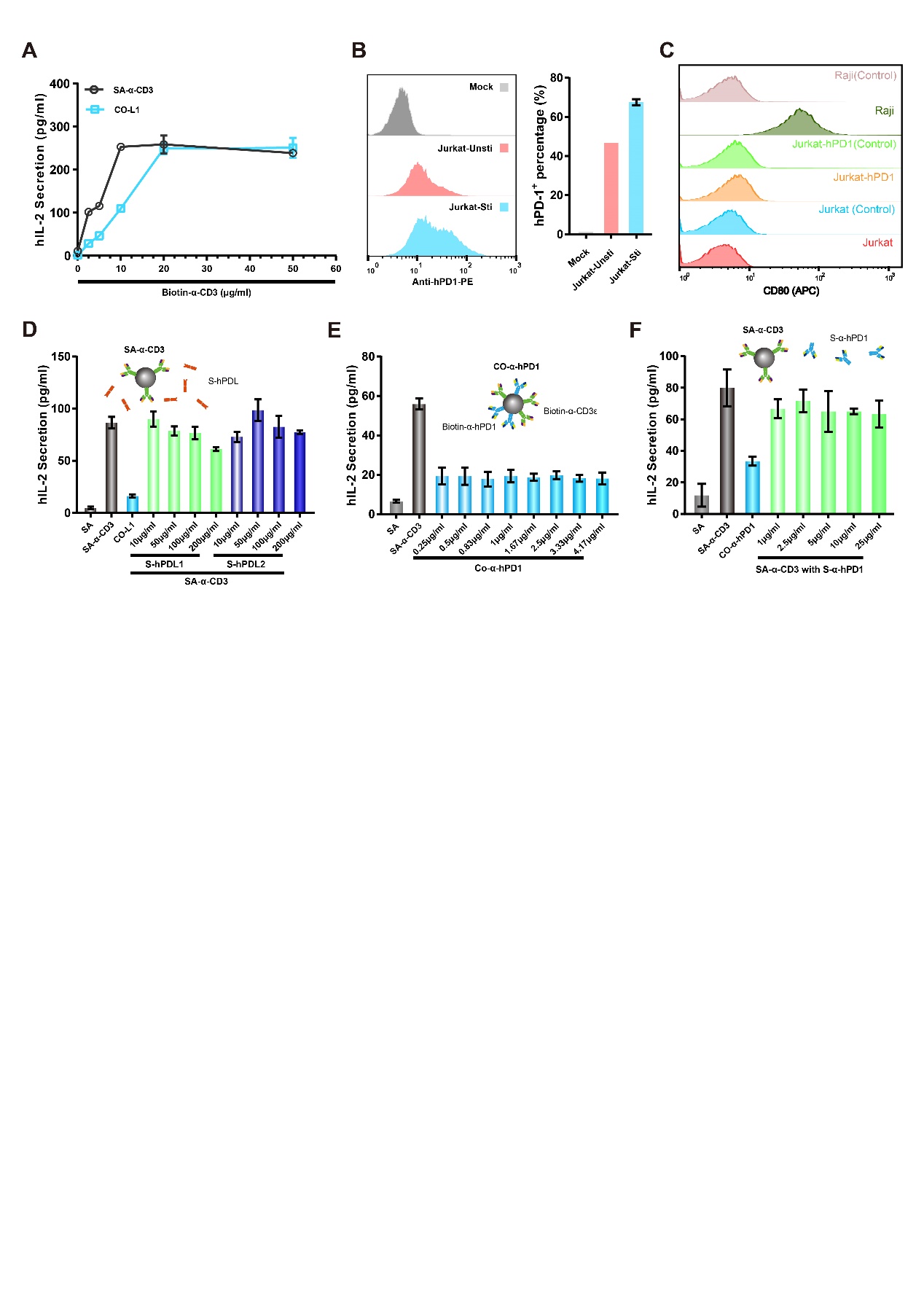

**Figure S5. Ligand immobilization is essential for PD-1 mediated T cell inhibition.**

**A**. IL-2 secretion of Jurkat cells cocultured with or without PD-L1 (15 μg/ml).

**B**. Cell surface expression of PD-1 in stimulated and unstimulated Jurkat cells.

**C**. The expression of CD80 in two Jurkat cell lines (Jurkat-wt and human PD-1 overexpressed Jurkat cell) and Raji cell.

**D-F**. IL-2 secretion of Jurkat cells stimulated with SA-α-CD3 beads in the presence of soluble PD-L1/PD-L2 (**D**), CO-α-hPD1 beads alone (**E**) or SA-α-CD3 beads with soluble PD-L1 antibody (**F**).

Note: In panels A, D, E, and F, 2.5μg/ml soluble anti-CD28 was used in the experiments, and all the experiments were repeated three times. IL-2 concentration of supernatant was quantified by ELISA kit.

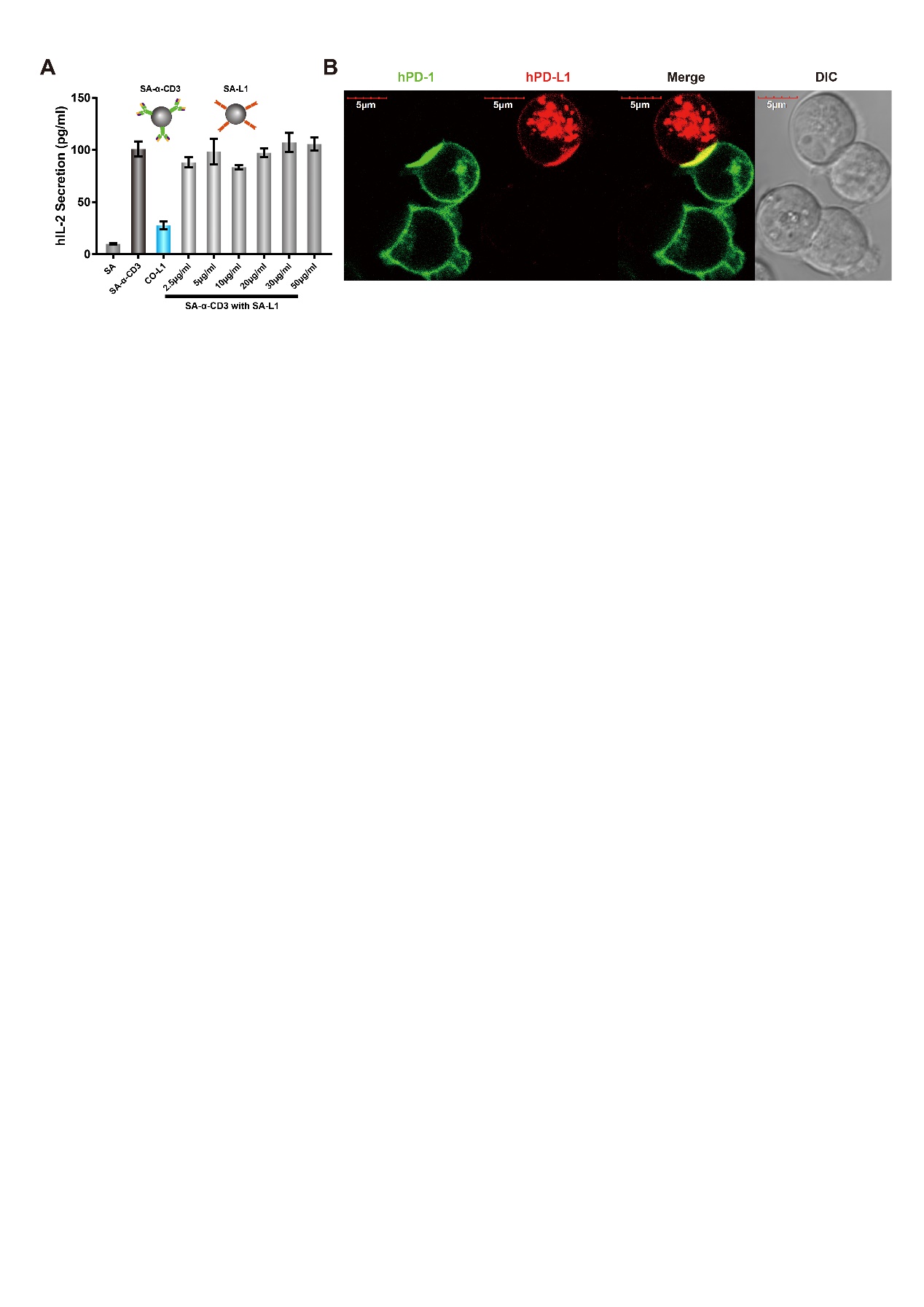

**Figure S6. PD-1 mediated inhibitory function depends on its IS localization.**

**A**. IL-2 secretion of Jurkat cells stimulated with anti-CD3 in the presence of co-immobilized PD-L1 (CO-L1) or PD-L1 immobilized on separated beads with indicated coating concentration.

**B**. Representative confocal image of Jurkat (hPD1^+^)-Raji (WT or hPD-L1^+^) conjugates. Cells were stained and imaged to visualize the interaction between Jurkat cells expressing human PD-1 (green) and Raji cells expressing PD-L1 (red).

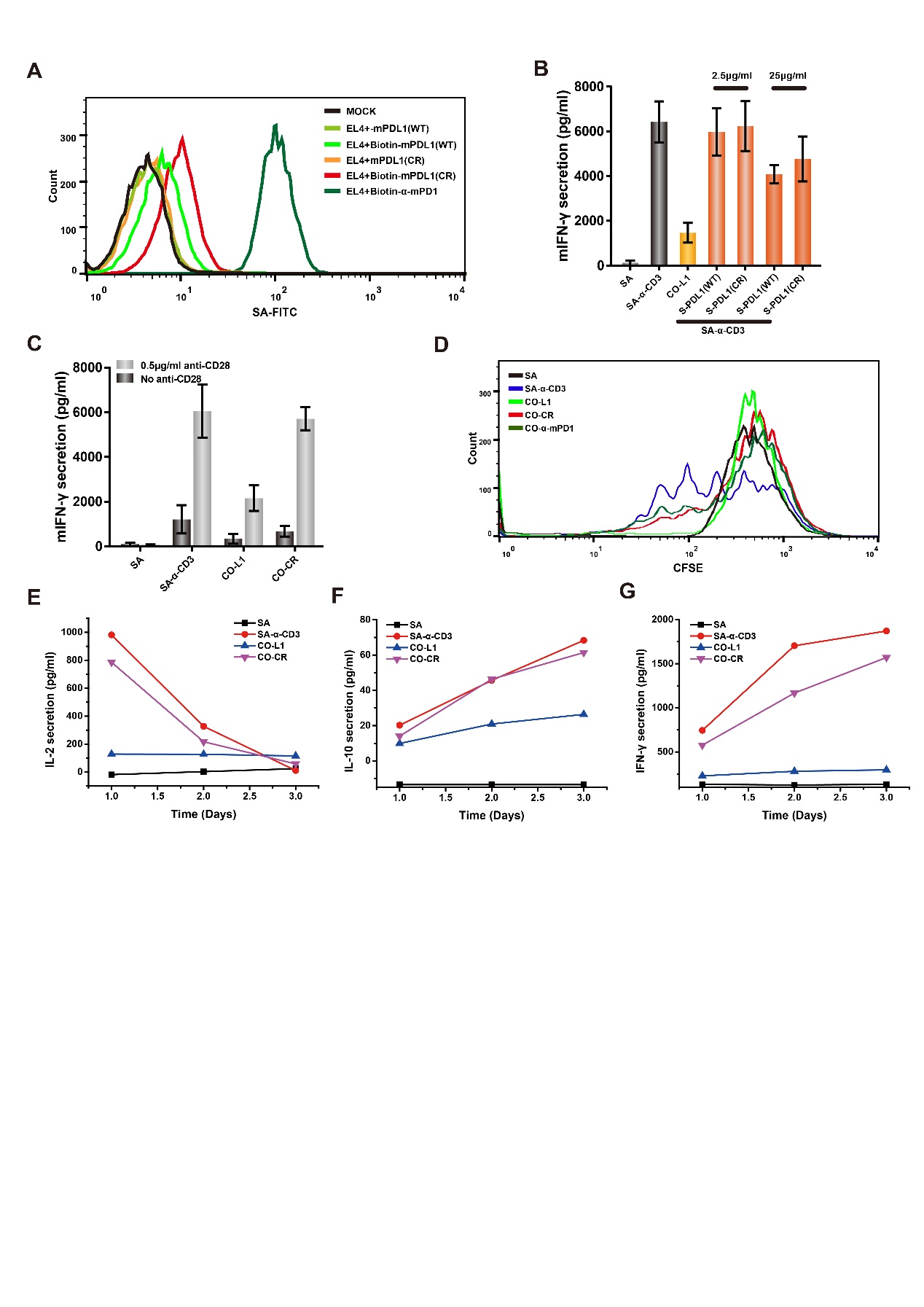

**Figure S7. Soluble mPD-L1-CR mutant block PD-1’s inhibitory function in mouse.**

**A**. Binding characteristics of wild type and CR mutated mouse PD-L1.

**B**. IFN-γ secretion of mouse primary CD8^+^ T cells cocultured with anti-CD3 beads in the presence of mouse S-mPDL1(WT) or mouse S-mPDL1(CR).

**C**. IFN-γ secretion of mouse primary CD8^+^ T cells stimulated with indicated beads in the presence or absence of mouse CD28 antibody (0.5μg/ml).

**D**. Proliferation of primary CD8^+^ T cells stimulated with indicated beads.

**E-G**. Cytokine secretion of mouse CD8^+^ T cells activated with indicated beads.
